# Metagenomic screening of global microbiomes identifies pathogen-enriched environments

**DOI:** 10.1101/376855

**Authors:** Xiaofang Li

## Abstract

**Background:** Human pathogens are widespread in the environment, and examination of pathogen-enriched environments in a rapid and high-throughput fashion is important for development of pathogen-risk precautionary measures.

**Methods:** In this study, a Local BLASTP procedure for metagenomic screening of pathogens in the environment was developed using a toxin-centered database. A total of 27 microbiomes derived from ocean water, freshwater, soil, feces, and wastewater were screened using the Local BLASTP procedure. Bioinformatic analysis and Canonical Correspondence Analysis were conducted to examine whether the toxins included in the database were taxonomically associated.

**Results:** The specificity of the Local BLASTP method was tested with known and unknown toxin sequences. Bioinformatic analysis indicated that most toxins were phylum-specific but not genus-specific. Canonical Correspondence Analysis implied that almost all of the toxins were associated with the phyla of *Proteobacteria*, *Nitrospirae* and *Firmicutes*. Local BLASTP screening of the global microbiomes showed that pore-forming RTX toxin and adenylate cyclase Cya were most prevalent globally in terms of relative abundance, while polluted water and feces samples were the most pathogen-enriched.

**Conclusions:** A Local BLASTP procedure was established for rapid detection of toxins in environmental samples. Screening of global microbiomes in this study provided a quantitative estimate of the most prevalent toxins and most pathogen-enriched environment.

## Introduction

Rapid identification of pathogens in a particular environment is important for pathogen-risk management. Human pathogens are ubiquitous in the environment, and infections from particular environments have been reported worldwide. For example, soil-related infectious diseases are common [1, 2]. *Legionella longbeachae* infection has been reported in many cases, mainly due to potting mixes and composts [3]. Survival of enteric viruses and bacteria has also been detected in various water environments, including aquifers and lakes [4-7].

Examination of pathogens from infected individuals with a particular clinical syndrome has been a major achievement of modern medical microbiology [8]. Nevertheless, we still know little about the magnitude of the abundance and diversity of known common pathogens in various environments, which is very important to the development of appropriate precautions for individuals who work or play with certain environmental substrates. This can be realized through metagenomic detection of pathogenic factors in a time-efficient and high-throughput manner using next-generation sequencing methods.

Metagenomic detection of pathogens can be accomplished through different schemes. Li et al. examined the level and diversity of bacterial pathogens in sewage treatment plants using a 16S rRNA amplicon-based metagenomic procedure [9]. Quantitative PCR has also been applied for monitoring specific pathogens in wastewater [10]. More studies have applied the whole-genome-assembly scheme to detect one or multiple dominant pathogens, most of which were for viral detection in clinical samples [11-14]. Although metagenomic-based whole-genome-assembly for bacterial pathogen detection can be conducted at the single species level [15], its computational requirements are high if in a high-throughput fashion. In 2014, Baldwin et al. [16] designed the PathoChip for screening pathogens in human tissues by targeting unique sequences of viral and prokaryotic genomes with multiple probes in a microarray. This approach can screen virtually all pathogen-enriched samples in a high-throughput manner.

Despite the aforementioned progress in metagenomic tools for pathogen detection, metagenomic screening for bacterial pathogens in environments such as soil, where microbial diversity is tremendous, is still challenging. This is mostly due to difficulty in assembling short reads generated by next-generation sequencing [8]. The whole-genome-assembly approach is efficient at identifying viromes, but not at dealing with bacterial communities. Amplicon-based approaches are able to detect bacterial pathogens in a high-throughput manner; however, it is well known that phenotypic diversity exists widely across and within microbial species of a genus because of divergent evolution [17, 18]. This also holds true for pathogenic factors [19]. Moreover, toxin factors, such as the Shiga toxin (*stx*) of *Shigella*, are primarily transferable through lateral gene transfer, which leads to the continuous evolution of pathogen species [20]. Therefore, it is necessary to examine the pathogen diversity in environmental metagenomes using essential virulence genes as biomarkers.

In this study, a toxin-centered virulence factors database was established, and the well-developed Local BLASTP method was applied to detect virulent factors in various environments globally. This procedure is metagenome-based and can be conducted in a high-throughput fashion, which greatly simplifies development of precautions for pathogen-enriched environments.

## Methods and Materials

### Environments and their metagenomes

Twenty-seven metagenomes were selected and downloaded from the MG-RAST server (Table 1). These metagenomes were derived from ocean water, freshwater, wastewater, natural soil, deserts and feces, representing the major environmental media found worldwide. Sequencing methods of the metagenomes include the Illumina, Ion Torrent and 454 platforms, and predicted proteins in the metagenomes ranged from 33,743 (fresh water, ID mgm4720261) to 1,966,121 (weedy garden soil, ID mgm4679254). The gene calling results were used for toxin factor screening in this study. The taxonomic composition at the genus level was also retrieved from the MG-RAST server for each metagenome.

**Table 1.**
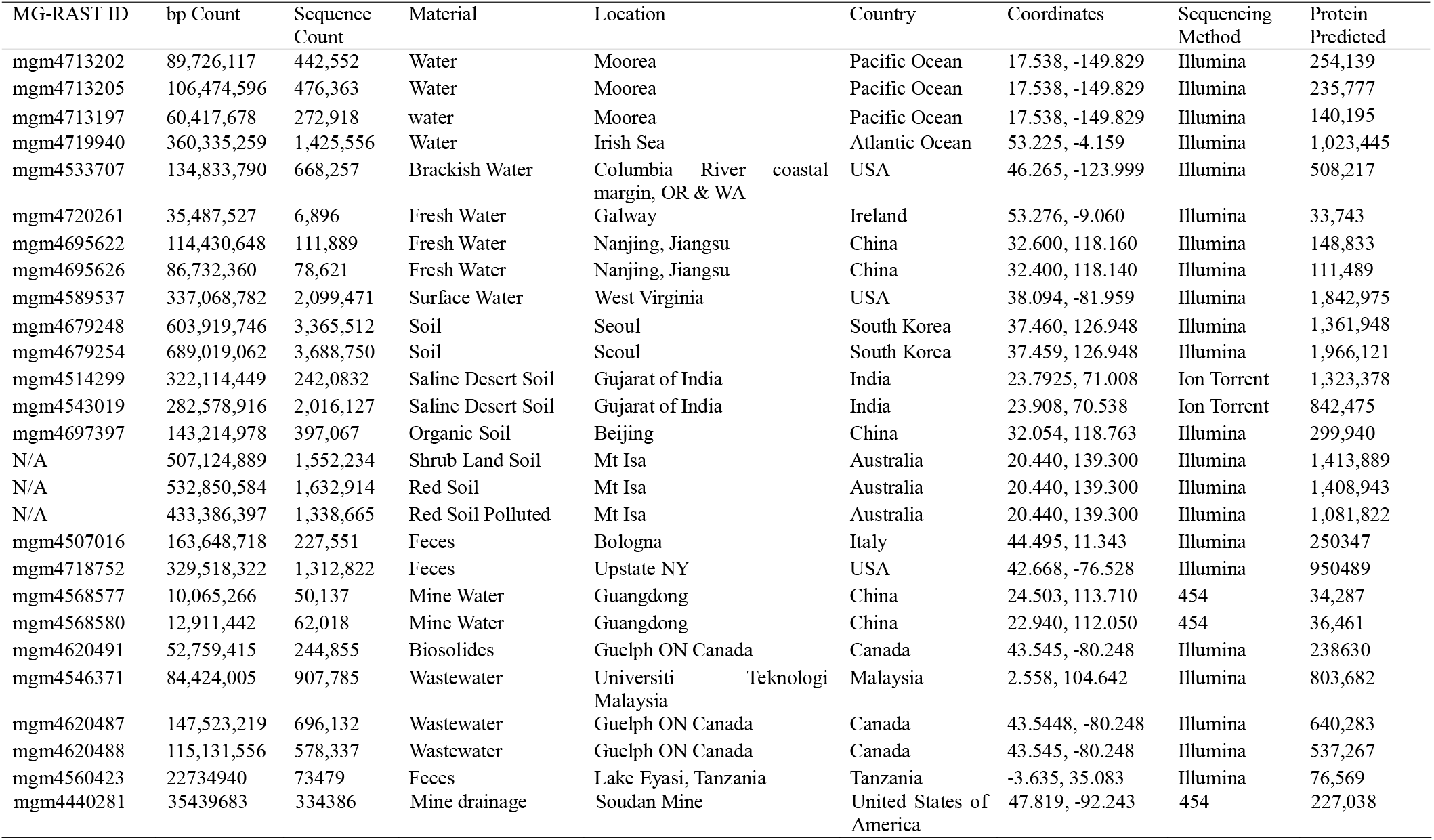
General information regarding the metagenomes retrieved from the MG-RAST server

### Toxin factor database

A toxin-centered database was established for bacterial pathogen detection in metagenomes in this study. Candidate toxin factors for pathogenic screening of environmental metagenomes were gathered based on well-studied pathogens summarized in Wikipedia^®^ under the entry of “pathogenic bacteria”, the Virulence Factor Database [21], a soil borne pathogen report by Jeffery and van der Putten [2],and a manure pathogen report by the United States Water Environment Federation [22]. Sequences of the toxin factors were then retrieved by searching the UniProt database using the toxin plus pathogen names as an entry [23], while typical homologs at a cutoff E value of 10^−6^ were gathered from GenBank based on BLAST results. Considering that virulence process involves several essential factors including toxins, various pathogen-derived secretion proteins were also included in the database, and it was tested that whether secretion proteins were as specific as toxin proteins for pathogen detection. The disease relevance of all virulence factors was screened using the WikiGenes system [24] and relevant publications (Table 2).

**Table 2.**
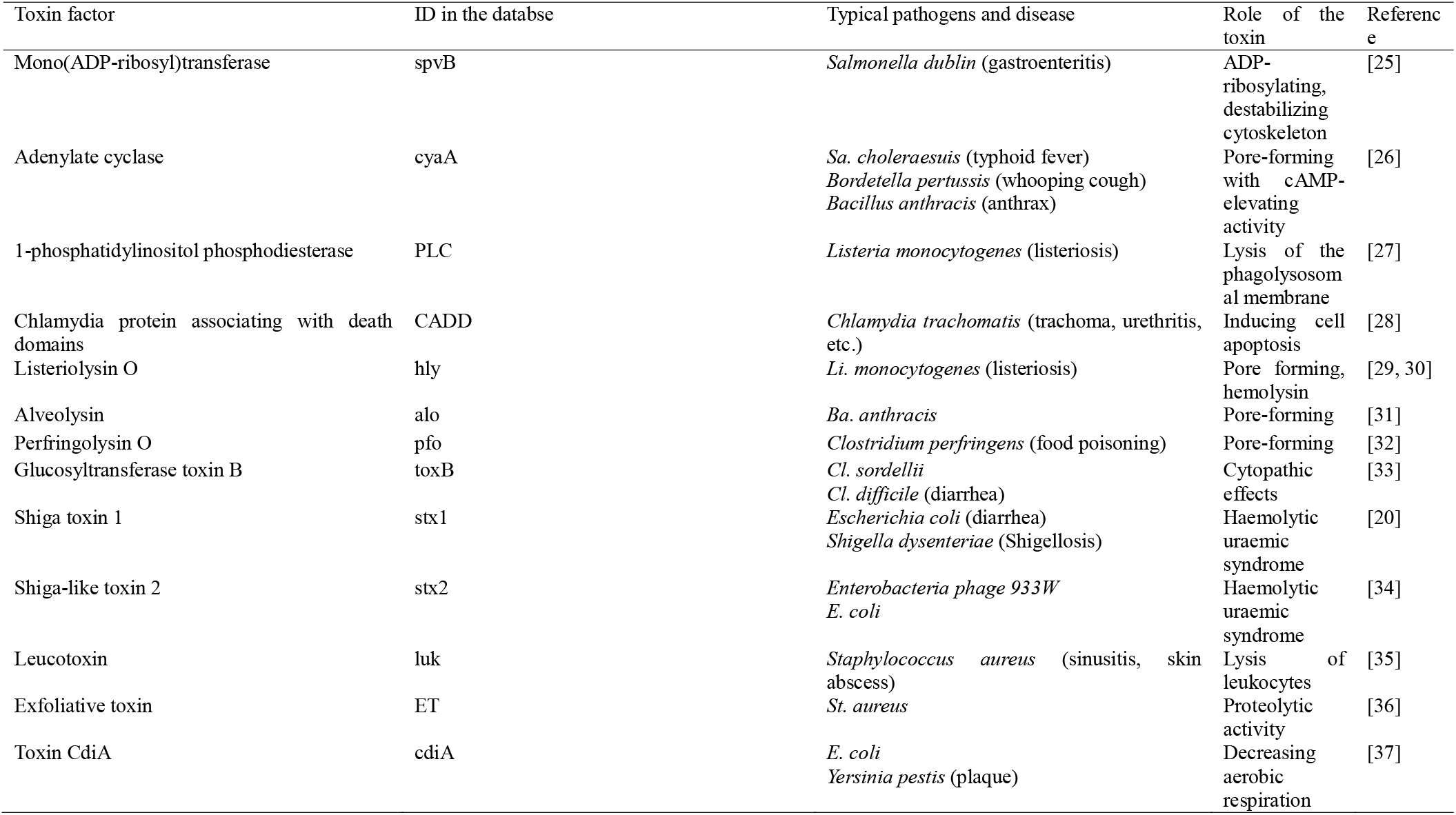
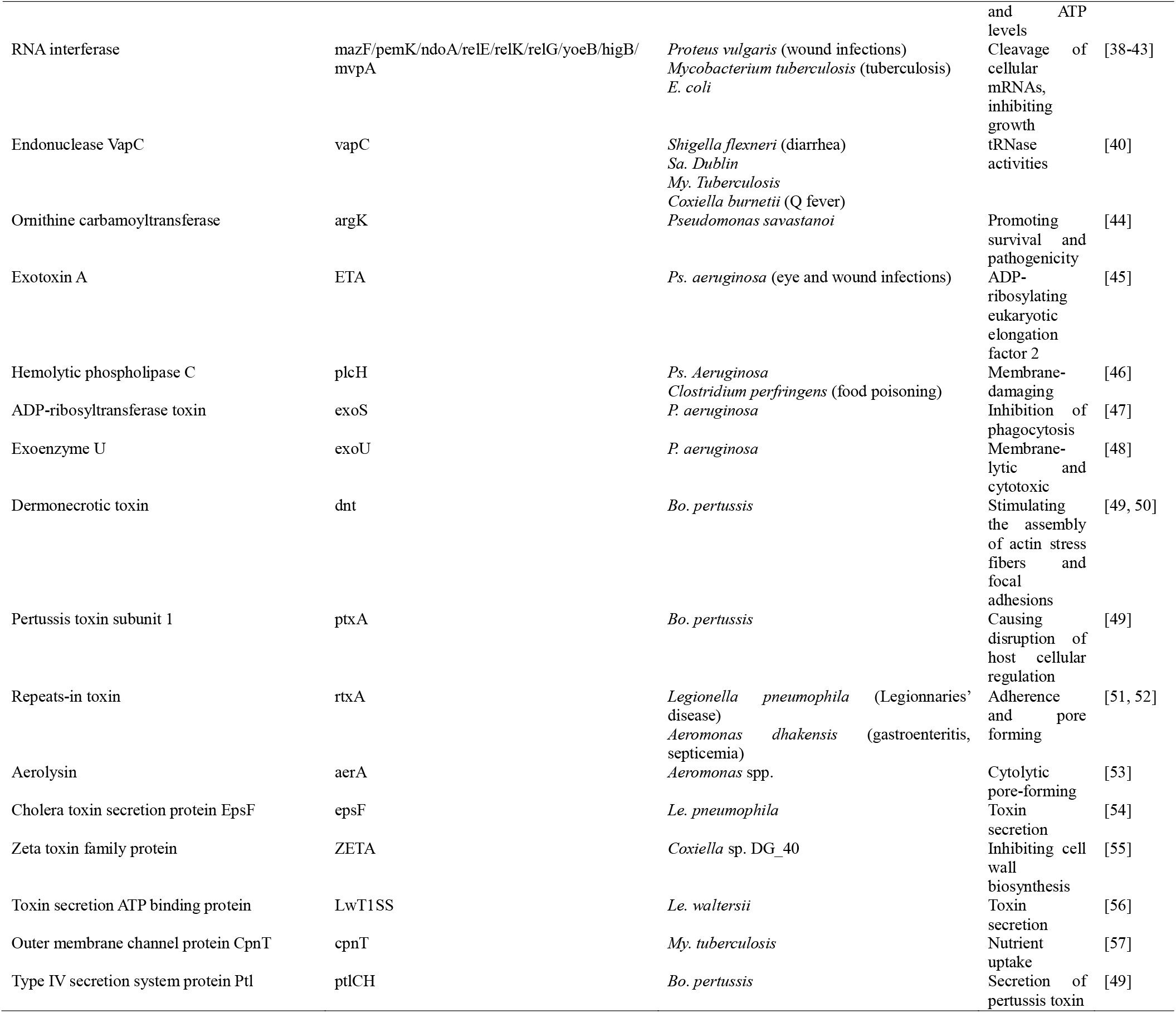
Typical virulence factors investigated in this study and their disease–relevance

### Local BLASTP

The Local BLASTP was applied following the procedure used in our previous study [58]. Basically, the gene calling results of each metagenome were searched against the toxin factor database using BLASTP embedded in BioEdit. The cutoff expectation E value was set as 10^−6^. The results of the Local BLASTP in BioEdit were then copied to an Excel worksheet, after which they were subjected to duplicate removal, quality control and subtotaled according to database ID. Duplicate removal was based on the hypothesis that each sequence contains one copy of a specific toxin factor, since the gene-calling results were used. For quality control of the BLSAT results, a cutoff value of 40% for amino acids identity and 20 aa (1/3 of the length of the shortest toxin factors (e.g., the Heat-Stable Enterotoxin C)) for query alignment length were used to filter the records. The toxins abundance matrix was formed for subsequent analyses.

### Specificity tests of the Local BLASTP method

Sequences from the toxin database established in this study, as “known sequences” to the database, were selected randomly and searched against the database using the BLASTP procedure. The genome of *Clostridium perfringens* ATCC 13124 (NC_008261), as “unknown” sequences to the database, was subject the Local BLASTX procedure as well. Homologous proteins were searched exhaustively in the GenBank database using BLASTP, with the representative toxin factors in the toxins database as a query. Sequences were retrieved and aligned using ClustalW, and Maximum-likelihood phylogeny was conducted with MEGA 7 [59].

### Data analysis

The toxin frequency in each metagenome was normalized to a total gene frequency of 1M to eliminate the effects of gene pool size. Toxin abundance in the 27 metagenomes was visualized using Circos [60]. The genus abundance of all metagenomes was calculated and sorted by genus name, followed by manual construction of a genus abundance matrix for subsequent biodiversity-toxin abundance Canonical Correspondence Analysis using R [61].

## Results and Discussion

In this study, a toxin-centered database was established for bacterial pathogen screening in various microbiomes globally through a Local BLASTP procedure. The specificity of the procedure was tested, the relative abundance of toxins in the microbiomes was examined, and the toxin-taxonomic abundance correspondence analysis was performed.

Like the previously established Local BLASTN method for antibiotic and metal resistance genes screening [58, 62, 63], the Local BLASTP method using the toxin-centered pathogen database in this study was successful at accurately identifying toxin proteins from the database. For screening of the *Clostridium perfringens* ATCC 13124 genome, the methods successfully detected the pore-forming genes and multiple copies of the glucosyltransferase (toxB-like) and ADP-ribosyltransferase (*spvB-like*) genes, based on the raw data. These results are consistent with the virulence genetic features of *Clostridium* sp. [21], which have not been well detailed in the GenBank annotation record. Such a cross-validation positively indicated that the Local BLASTP procedure established here is useful in predicting toxin genes in unknown genomes. Yet for a semi-quantitative method to estimate toxin factors in metagenomes, a false positive analysis is required to examine to what level mismatch is included in the Local BLASTP results. Actually, the cutoff values of identity greatly impact the homolog virulence factor abundance returned. At cutoff values of 40% for identify and 20 aa for alignment length, only four records for *Clostridium perfringens* ATCC 13124 genome query were returned after duplication removal, one for 1-phosphatidylinositol phosphodiesterase, one for pore-forming alveolysin, one for Ornithine carbamoyltransferase and one for RNA interferase NdoA. At a cutoff identity value of 35%, one more record (Toxin secretion ATP binding protein) was returned. This means that the Local BLASTP procedure was able to detect the virulence factors in unknown genomic dataset at least semi-quantitatively, with proper cutoff values for data quality control. The accuracy of the BLASTP procedure in virulence factor detection was further tested using the genomes of *Bacillus thuringiensis serovar konkukian* str. 97-27 (AE017355.1) and Helicobacter pylori 26695 (AE000511.1) (results not shown).

As mentioned above, functional genes including toxin factors may partly evolve through lateral gene transfer, which makes their taxonomic affiliation difficult. It is thus interesting to explore how specific toxin factors are associated with the taxonomic units of pathogens. Here, I explored this issue by investigating the taxonomic distribution of homologs of toxinsretrieved from the GenBank database. Generally, at a lower expectation value, most toxins were associated with a specific group of pathogens. For example, at a cutoff E value of 10^−6^ (the default unless specified), 241 out of 242 returned records of *Mycobacterium tuberculosis* RelEhomologs fell within the phylum *Actinobacteria*. Moreover, 89% of these homologs were from the genus *Mycobacterium*, while 99.7% of *Yersinia pestis* CdiAhomologs and 92.7% of *Bordetella pertussis* cya homologs belonged to *Proteobacteria*, and homologs of *Aeromonas dhakensis* repeats-in toxin (RtxA) were mostly associated with the class *Gammaproteobacteria* (206 out of 242). However, no obvious genus-toxin association was identified. It is worth noting that these results largely depended on the availability of toxin sequences in each taxonomic unit. The lack of a genus-toxin association basically denied the possibility of detecting a specific pathogen using a specific toxin as a single signature [16].

It is still not clear whether virulence secretion proteins are specific for pathogen detection as signatures, through they are essential for virulence process [20]. For example, the contact-dependent toxin delivery protein CdiA was found to be widespread in bacteria [37]. The relative abundance of secretion proteins in the 27 microbiomes was determined as well as that of the toxins which are essential to virulence processes. The results of the present study showed that the abundance of secretion proteins selected in the database was strongly correlated with the toxin abundance (R^2^ = 0.80, Figure 1). The most abundant secretion proteins included *L. waltersii* toxin secretion protein (LWT1SS), *L. pneumophila* toxin secretion protein ApxIB, and *Aeromonas hydrophila* RTX transporter (RtxB) (data not shown). Further exploration indicated that although *A. hydrophila* RtxB homologs from GenBank were found in all *Proteobacteria* classes, most of the RtxB-harboring species have been reported to be pathogens, including *Vibrio* spp. [64], *Pseudomonas* spp., *Neisseria meningitides* [65], *Ralstonia* spp. [66], and *Yersinia* spp. [21]. This may imply the pathogen-specific nature of secretion proteins included in the database, and that toxin secretion proteins can be used as signatures for pathogen detection as well.

**Figure 1.**
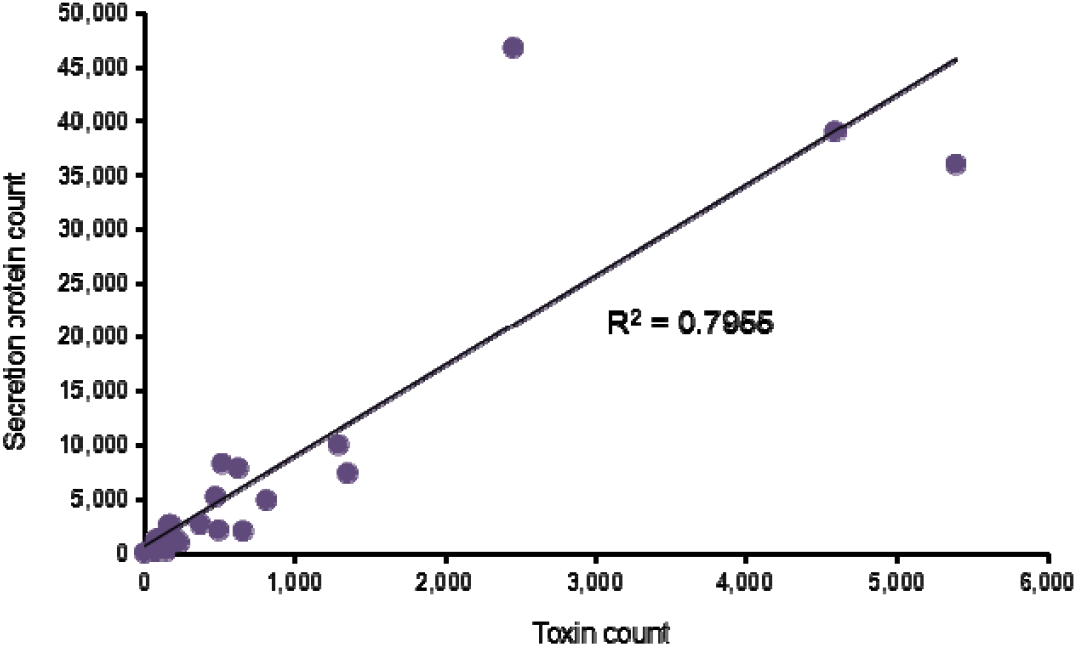
Correlation between relative abundance of toxins and secretion proteins in the global microbiomes.

Toxin-phyla CCA results showed that all phyla can be clearly separated into two groups, and that almost all toxins were associated with *Proteobacteria, Nitrospirae* and *Firmicutes* (Figure 2). Considering the phylum-specificity of the toxins stated above, these results can be biased because of the taxonomic affiliation of toxins included in the Local BLASTP database. The taxonomic distribution proportion of currently available genomes of identified pathogens was reflected in the toxin database, with *Proteobacteria* and *Firmicutes* accounting for the majority of the genomes. However, the CCA results may also indicate, at least in part, a proportional lack of pathogens in some phyla, such as *Crenarchaeota, Euryarchaeota, Verrucomicrobia* and *Bacteroidetes* [67]. Archaea cannot easily absorb phage particles because of their extracellular structures, which differ from bacteria [68]. A recent study by Li et al. also found that the five most abundant bacterial pathogens were from either *Proteobacteria* or *Firmicutes* in wastewater microbiomes [9]. Taken together, these findings could indicate that *Proteobacteria* or *Firmicutes* were evolutionarily enriched with pathogens when they dominated most environmental microbiomes on the planet [69, 70].

**Figure 2.**
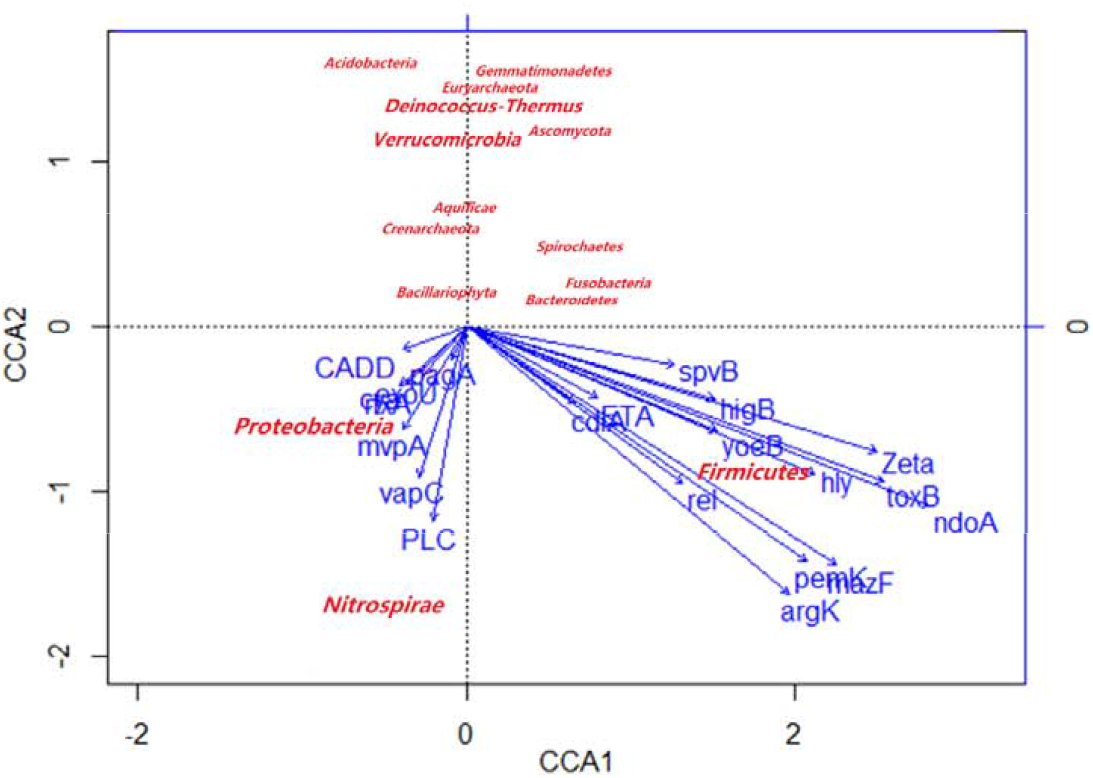
Canonical correspondence analysis of the associations between phyla and toxins.

Interestingly, there was a strong association between the phylum *Nitrospirae* and toxins of RNase inteferases (MvpA and MapC) and *Listeria monocytogenes* 1-phosphatidylinositol phosphodiesterase PLC.

Further searches against the UniProt database [71] revealed no homologous records of MvpA and PLC from *Nitrospirae*, and only 109 out of 15,574 bacterial records for VapC were from *Nitrospirae*. These findings imply that there are many more *Nitrospirae* pathogens harboring MvpA and PLC that have yet to be discovered.

The screening of toxins in the 27 global microbiomes revealed the most prevalent toxins and pathogen-enriched environment. Specifically, the results showed that the RTX toxin RtxA and adenylate cyclase Cya were most prevalent globally in terms of relative abundance. RTX toxins comprise a large family of pore-forming exotoxins. Known homologs in the GenBank database of *Aeromonas dhakensis* RtxA were mainly in the genera of *Aeromonas, Pseudomonas* (e.g., CP015992), *Vibrio* (e.g., CP002556) and *Legionella* (e.g., CP015953). These genera are well known to be associated with gastroenteritis, eye and wound infections, cholera and legionellosis, and RTX toxins are a key part of the virulence systems of each of these conditions [72-75]. Cya is an essential unit of *Bacillus* anthracisvirulence that causes anthrax and may lead to mammalian death [76]. Known homologs in the GenBank database of *Bacillus anthracis* Cya were mainly from *Bacillus* spp., *Bordetella* spp., *Pseudomonas aeruginosa, Yersiniapseudotuberculosis*, and *Vibrio* spp. Their presence in the environment should be carefully examined and precautions should be taken to prevent infection by these organisms since many of them are associated with very common diseases such as whooping cough.

The main purpose of the Local BLASTP method established here was to screen pathogen-enriched environments to enable development of precautionary measures. Our results clearly indicated that contaminated lake water, feces and wastewater microbiomes were rich in pathogens (Figure 3). Although there was no detailed background information regarding these environments in this study, the results presented herein may provide important implications for pathogen-related risk control. Surprisingly, two lake water microbiomes from Nanjing, China contained the highest toxin factors among the 27 samples. Further investigation of the location and contamination status supported the sewage-nature of the lake water. In China, most polluted lakes receive sewage that includes feces materials [77]. According to an official survey conducted in 2015, Nanjing has 28 lakes with a total area of 14 km^2^, among which 96.7% are classified as polluted (Class V of the national standard). Studies have documented that pathogens tend to be enriched in polluted waters [14]. It is not surprising to find that feces samples had very high abundance of toxins. Epidemical statistics have indicated that feces are the most important pathway for diarrheal diseases, which is a leading cause of childhood death globally [78]. Thus, the present study provides a method for obtaining quantitative estimates of pathogen enrichment of various environments, and polluted freshwater systems are found to be highly pathogen-enriched relative to safer environments such as ocean water and natural soils.

**Figure 3.**
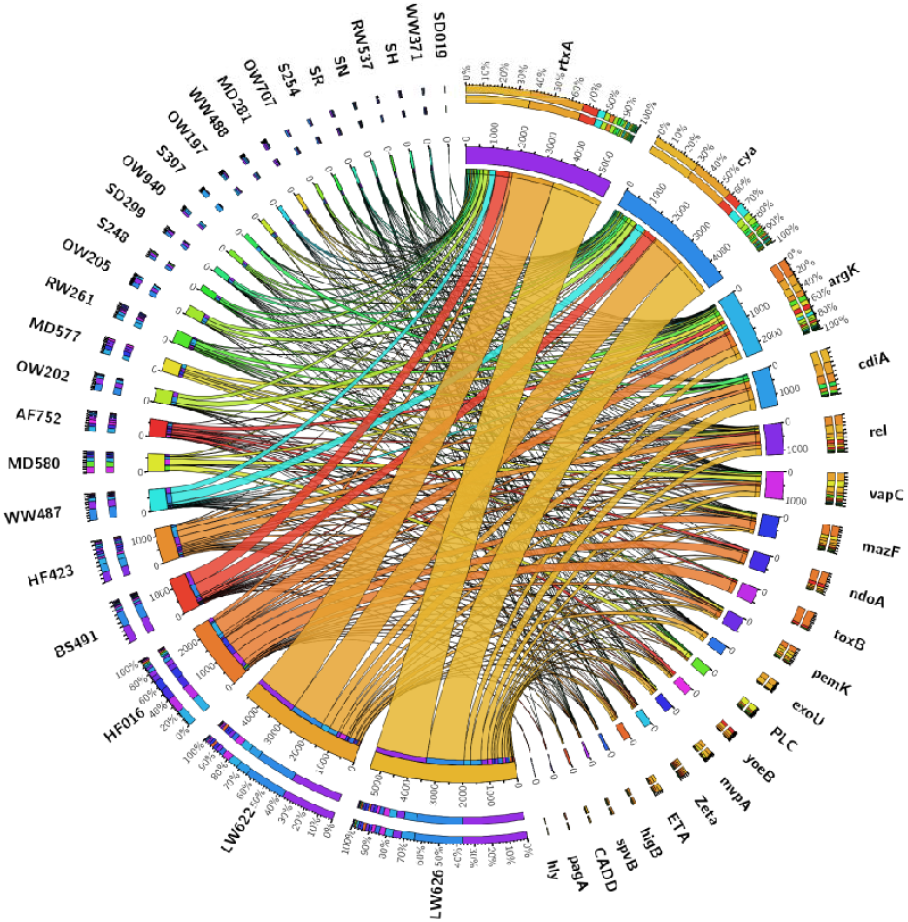
Circular visualization of the toxin abundance in the microbiomes selected from locations worldwide. The designated environment was prefixed with the first letters of the environment names and suffixed with the last three numbers of their MG-RAST ID in Table 1.

## Conclusions

A Local BLASTP procedure was established for rapid detection of toxins in environmental samples. Screening of global microbiomes in this study provided a quantitative estimate of the most prevalent toxins and most pathogen-enriched environments.

## Declarations

### Acknowledgement

I thank Dr. Philip L. Bond and The University of Queensland for providing training in bioinformatics. We would like to thank LetPub (www.letpub.com) for providing linguistic assistance during the preparation of this manuscript. I also thank the founders of the existing pathogen-relevant database, particularly the Virulence Factor Database, which provided valuable reference for the buildup of the toxin database in this study.

### Availability of data and materials

The toxin database is available in the Supplementary Materials. All toxin abundance data in this study can be provided by the author upon request.

### Authors’ contributions

XL initiated the study, analyzed the data and wrote the manuscript.

### Ethics approval

Not applicable.

### Consent for publication

Not applicable.

## References

[1] Baumgardner DJ. Soil-related bacterial and fungal infections. J Am Board Fam Med, 2012, 25: 734–744

[2] Jeffery S, van der Putten WH, Ispra, Italy: Joint Research Centre, Institute for Environment and Sustainability 2011

[3] Whiley H, Bentham R. *Legionella longbeachae* and legionellosis. Emerging Infectious Diseases, 2011, 17: 579–583

[4] Wait DA, Sobsey MD. Comparative survival of enteric viruses and bacteria in Atlanticocean seawater. Water Sci Technol, 2001, 43: 139–142

[5] Dan TBB, Wynne D, Manor Y. Survival of enteric bacteria and viruses in lake kinneret, Israel. Water Res, 1997, 31: 2755–2760

[6] Keswick BH, Gerba CP, Secor SL, et al. Survival of enteric viruses and indicator bacteria in groundwater. J Environ Sci Heal A, 1982, 17: 903–912

[7] Cooper RC, Golueke CG. Survival of enteric bacteria and viruses in compost and its leachate. Compost Sci Land Ut, 1979, 20: 29–35

[8] Miller RR, Montoya V, Gardy JL, et al. Metagenomics for pathogen detection in public health. Genome Med, 2013, 5:

[9] Li B, Ju F, Cai L, et al. Profile and fate of bacterial pathogens in sewage treatment plants revealed by high-throughput metagenomic approach. Environ Sci Technol, 2015, 49: 10492–10502

[10] Amha YM, Anwar MZ, Kumaraswamy R, et al. *Mycobacteria* in municipal wastewater treatment and reuse: Microbial diversity for screening the occurrence of clinically and environmentally relevant species in arid regions. Environ Sci Technol, 2017, 51: 3048–3056

[11] Yang J, Yang F, Ren LL, et al. Unbiased parallel detection of viral pathogens in clinical samples by use of a metagenomic approach. J Clin Microbiol, 2011, 49: 3463–3469

[12] Granberg F, Vicente-Rubiano M, Rubio-Guerri C, et al. Metagenomic detection of viral pathogens in spanish honeybees: Co-infection by aphid lethal paralysis, Israel acute paralysis and lake sinai viruses. Plos One, 2013, 8:

[13] Nakamura S, Yang CS, Sakon N, et al. Direct metagenomic detection of viral pathogens in nasal and fecal specimens using an unbiased high-throughput sequencing approach. Plos One, 2009, 4: e4219

[14] Bibby K. Metagenomic identification of viral pathogens. Trends Biotechnol, 2013, 31: 11–15

[15] Fukui Y, Aoki K, Okuma S, et al. Metagenomic analysis for detecting pathogens in culture-negative infective endocarditis. J Infect Chemother, 2015, 21: 882–884

[16] Baldwin DA, Feldman M, Alwine JC, et al. Metagenomic assay for identification of microbial pathogens in tumor tissues. Mbio, 2014, 5: 01714–14

[17] Achtman M, Wagner M. Microbial diversity and the genetic nature of microbial species. Nat Rev Micro, 2008, 6: 431–440

[18] Liao PY, Lee KH. From snps to functional polymorphism: The insight into biotechnology applications. Biochem Eng J, 2010, 49: 149–158

[19] Sokurenko EV, Hasty DL, Dykhuizen DE. Pathoadaptive mutations: Gene loss and variation in bacterial pathogens. Trends Microbiol, 1999, 7: 191–195

[20] Strauch E, Lurz R, Beutin L. Characterization of a shiga toxin-encoding temperate bacteriophage of *Shigella sonnei.* Infect Immun, 2001, 69: 7588–7595

[21] Chen LH, Zheng DD, Liu B, et al. Vfdb 2016: Hierarchical and refined dataset for big data analysis-10 years on. Nucleic Acids Res, 2016, 44: D694–D697

[22] US Water Environment Federation. Manure pathogens: Manure management, regulations, and water quality protection. Water Environment Federation Alexandria, Virginia: WEF Press, 2009

[23] Bateman A, Martin MJ, O’Donovan C, et al. Uniprot: The universal protein knowledgebase. Nucleic Acids Res, 2017, 45: D158–D169

[24] Hoffmann R. A wiki for the life sciences where authorship matters. Nature Genetics, 2008, 40: 1047–1051

[25] Lesnick ML, Reiner NE, Fierer J, et al. The *SalmonellaspvB* virulence gene encodes an enzyme that ADP-ribosylates actin and destabilizes the cytoskeleton of eukaryotic cells. Mol Microbiol, 2001, 39: 1464–1470

[26] Skopova K, Tomalova B, Kanchev I, et al. Camp-elevating capacity of the adenylate cyclase toxin-hemolysin is sufficient for lung infection but not for full virulence of bordetella pertussis. Infect Immun, 2017,

[27] Henner DJ, Yang M, Chen E, et al. Sequence of the *Bacillus thuringiensis* phosphatidylinositol specific phospholipase-c. Nucleic Acids Res, 1988, 16: 10383–10383

[28] Schwarzenbacher R, Stenner-Liewen F, Liewen H, et al. Structure of the chlamydia protein cadd reveals a redox enzyme that modulates host cell apoptosis. J Biol Chem, 2004, 279: 29320–29324

[29] Hamon MA, Batsche E, Regnault B, et al. Histone modifications induced by a family of bacterial toxins. P Natl Acad Sci USA, 2007, 104: 13467–13472

[30] Cossart P. The listeriolysin o-gene - a chromosomal locus crucial for the virulence of *Listeria monocytogenes*. Infection, 1988, 16: S157–S159

[31] Geoffroy C, Mengaud J, Alouf JE, et al. Alveolysin, the thiol-activated toxin of *Bacillus alvei*, is homologous to listeriolysin o, perfringolysin o, pneumolysin, and streptolysin o and contains a single cysteine. J Bacteriol, 1990, 172: 7301–7305

[32] Rossjohn J, Polekhina G, Feil SC, et al. Structures of perfringolysin o suggest a pathway for activation of cholesterol-dependent cytolysins. J Mol Biol, 2007, 367: 1227–1236

[33] Lyras D, O’Connor JR, Howarth PM, et al. Toxin b is essential for virulence of *Clostridium difficile.* Nature, 2009, 458: 1176–1181

[34] Schmidt H, Scheef J, JanetzkiMittmann C, et al. An ilex trna gene is located close to the shiga toxin ii operon in enterohemorrhagic *Escherichia coli* o157 and non-o157 strains. FEMS Microbiol Lett, 1997, 149: 39–44

[35] Labandeira-Rey M, Couzon F, Boisset S, et al. *Staphylococcus aureus* panton-valentine leukocidin causes necrotizing pneumonia. Science, 2007, 315: 1130–1133

[36] Bukowski M, Wladyka B, Dubin G. Exfoliative toxins of *Staphylococcus aureus.* Toxins, 2010, 2: 1148–1165

[37] Aoki SK, Diner EJ, de Roodenbeke CT, et al. A widespread family of polymorphic contact-dependent toxin delivery systems in bacteria. Nature, 2010, 468: 439–442

[38] Tian QB, Ohnishi M, Tabuchi A, et al. A new plasmid-encoded proteic killer gene system: Cloning, sequencing, and analysing *hig* locus of plasmid rts1. Biochem Bioph Res Co, 1996, 220: 280–284

[39] Hurley JM, Woychik NA. Bacterial toxin HigB associates with ribosomes and mediates translation-dependent mRNA cleavage at a-rich sites. J Biol Chem, 2009, 284: 18605–18613

[40] Pullinger GD, Lax AJ. A Salmonella-dublin virulence plasmid locus that affects bacterial-growth under nutrient-limited conditions. Mol Microbiol, 1992, 6: 1631–1643

[41] Korch SB, Contreras H, Clark-Curtiss JE. Three *Mycobacterium tuberculosisRel* toxin-antitoxin modules inhibit mycobacterial growth and are expressed in infected human macrophages. J Bacteriol, 2009, 191: 1618–1630

[42] Pellegrini O, Mathy N, Gogos A, et al. *The Bacillus* subtilis ydcde operon encodes an endoribonuclease of the *mazF/pemK* family and its inhibitor. Molecular Microbiology, 2005, 56: 1139–1148

[43] Yamaguchi Y, Inouye M. Regulation of growth and death in *Escherichia coli* by toxin-antitoxin systems. Nature Rev Microbiol, 2011, 9: 779–790

[44] Hatziloukas E, Panopoulos NJ. Origin, structure, and regulation of argk, encoding the phaseolotoxin-resistant ornithine carbamoyltransferase in pseudomonas syringae pv. Phaseolicola, and functional expression of argk in transgenic tobacco. J Bacteriol, 1992, 174: 5895–5909

[45] Yates SP, Merrill AR. Elucidation of eukaryotic elongation factor-2 contact sites within the catalytic domain of pseudomonas aeruginosa exotoxin a. Biochem J, 2004, 379: 563–572

[46] Songer JG. Bacterial phospholipases and their role in virulence. Trends Microbiol, 5: 156–161

[47] Krueger KM, Barbieri JT. The family of bacterial ADP-ribosylating exotoxins. Clin Microbiol Rev, 1995, 8: 34–47

[48] Phillips RM, Six DA, Dennis EA, et al. In vivo phospholipase activity of the *Pseudomonas* aeruginosa cytotoxin *exoUand* protection of mammalian cells with phospholipase a2 inhibitors. J Biol Chem, 2003, 278: 41326–41332

[49] Weiss AA, Johnson FD, Burns DL. Molecular characterization of an operon required for pertussis toxin secretion. P Natl Acad Sci USA, 1993, 90: 2970–2974

[50] Masuzawa T, Sawaki K, Nagaoka H, et al. Relationship between pathogenicity of coxiella burnetii isolates and gene sequences of the macrophage infectivity potentiator (cbmip) and sensor-like protein (qrsa). FEMS Microbiol Lett, 1997, 154: 201–205

[51] D’Auria G, Jimenez N, Peris-Bondia F, et al. Virulence factor rtx in *Legionella pneumophila*, evidence suggesting it is a modular multifunctional protein. Bmc Genomics, 2008, 9:

[52] Rasmussen-Ivey CR, Figueras MJ, McGarey D, et al. Virulence factors of *Aeromonas hydrophila:* In the wake of reclassification. Frontiers in Microbiology, 2016, 7:

[53] Howard SP, Garland WJ, Green MJ, et al. Nucleotide sequence of the gene for the hole-forming toxin aerolysin of aeromonas hydrophila. J Bacteriol, 1987, 169: 2869–2871

[54] Sandkvist M, Michel LO, Hough LP, et al. General secretion pathway (eps) genes required for toxin secretion and outer membrane biogenesis in vibrio cholerae. J Bacteriol, 1997, 179: 6994–7003

[55] Lioy VS, Machon C, Tabone M, et al. The ζ toxin induces a set of protective responses and dormancy. Plos One, 2012, 7: e30282

[56] Söderberg MA, Rossier O, Cianciotto NP. The type ii protein secretion system of legionella pneumophila promotes growth at low temperatures. J Bacteriol, 2004, 186: 3712–3720

[57] Danilchanka O, Pires D, Anes E, et al. The mycobacterium tuberculosis outer membrane channel protein cpnt confers susceptibility to toxic molecules. Antimicrob Agents Ch, 2015, 59: 2328–2336

[58] Li X, Zhu YG, Shaban B, et al. Assessing the genetic diversity of cu resistance in mine tailings through high-throughput recovery of full-length copa genes. Sci Rep, 2015, 5: 13258

[59] Kumar S, Stecher G, Tamura K. Mega7: Molecular evolutionary genetics analysis version 7.0 for bigger datasets. Mol Biol Evol, 2016, 33: 1870–1874

[60] Krzywinski M, Schein J, Birol I, et al. Circos: An information aesthetic for comparative genomics. Genome Res, 2009, 19: 1639–1645

[61] R Core Team. R: A language and environment for statistical computing. 2016, URL http://www.R-project.org/.

[62] Li XF, Bond PL, Huang LB. Diversity of as metabolism functional genes in Pb-Zn mine tailings. Pedosphere, 2017, 27: 630–637

[63] Gupta SK, Padmanabhan BR, Diene SM, et al. Arg-annot, a new bioinformatic tool to discover antibiotic resistance genes in bacterial genomes. Antimicrob Agents Ch, 2014, 58: 212–220

[64] Austin B, Zhang XH. Vibrio harveyi: A significant pathogen of marine vertebrates and invertebrates. Lett Appl Microbiol, 2006, 43: 119–124

[65] Rouphael NG, Stephens DS. Neisseria meningitidis: Biology, microbiology, and epidemiology. Methods in molecular biology (Clifton, NJ), 2012, 799: 1–20

[66] Xu J, Zheng HJ, Liu L, et al. Complete genome sequence of the plant pathogen ralstonia solanacearum strain po82. J Bacteriol, 2011, 193: 4261–4262

[67] Ecker DJ, Sampath R, Willett P, et al. The microbial rosetta stone database: A compilation of global and emerging infectious microorganisms and bioterrorist threat agents. Bmc Microbiol, 2005, 5:

[68] Gill EE, Brinkman FSL. The proportional lack of archaeal pathogens: Do viruses/phages hold the key? Bioessays, 2011, 33: 248–254

[69] Roesch LF, Fulthorpe RR, Riva A, et al. Pyrosequencing enumerates and contrasts soil microbial diversity. Isme J, 2007, 1: 283–290

[70] Fierer N, Bradford MA, Jackson RB. Toward an ecological classification of soil bacteria. Ecology, 2007, 88: 1354–1364

[71] Apweiler R, Bairoch A, Wu CH, et al. Uniprot: The universal protein knowledgebase. Nucleic Acids Res, 2004, 32: D115–D119

[72] Lin W, Fullner KJ, Clayton R, et al. Identification of a vibrio cholerae rtx toxin gene cluster that is tightly linked to the cholera toxin prophage. P Natl Acad Sci USA, 1999, 96: 1071–1076

[73] Cirillo SLG, Bermudez LE, El-Etr SH, et al. Legionella pneumophila entry gene *rtxa* is involved in virulence. Infect Immun, 2001, 69: 508–517

[74] Suarez G, Khajanchi BK, Sierra JC, et al. Actin cross-linking domain of aeromonas hydrophila repeat in toxin a (*rtxa*) induces host cell rounding and apoptosis. Gene, 2012, 506: 369–376

[75] Terada LS, Johansen KA, Nowbar S, et al. Pseudomonas aeruginosa hemolytic phospholipase c suppresses neutrophil respiratory burst activity. Infect Immun, 1999, 67: 2371–2376

[76] Leppla SH. Anthrax toxin edema factor - a bacterial adenylate-cyclase that increases cyclic-amp concentrations in eukaryotic cells. P Natl Acad Sci-Biol, 1982, 79: 3162–3166

[77] Qiu Z. Current pollution status of china’s lakes. The 5th Forum for China Lakes, 2015, 5.

[78] Liu L, Johnson HL, Cousens S, et al. Global, regional, and national causes of child mortality: An updated systematic analysis for 2010 with time trends since 2000. Lancet, 2012, 379: 2151–2161

